# Learning sequence patterns of AGO-sRNA affinity from high-throughput sequencing libraries to improve *in silico* functional small RNA detection and classification in plants

**DOI:** 10.1101/173575

**Authors:** Lionel Morgado, Ritsert C. Jansen, Frank Johannes

**Affiliations:** Groningen Bioinformatics Centre, University of Groningen, Groningen, 9747 AG Groningen, The Netherlands; Department of Plant Sciences, Technical University of Munich, Freising, 85354 Freising, Germany; Institute for Advanced Study, Technical University of Munich, Garching, 85748 Garching, Germany

## Abstract

The loading of small RNA (sRNA) into Argonaute (AGO) complexes is a crucial step in all regulatory pathways identified so far in plants that depend on such non-coding sequences. Important transcriptional and post-transcriptional silencing mechanisms can be activated depending on the specific AGO protein to which sRNA bind. It is known that sRNA-AGO associations are at least partly encoded in the sRNA primary structure, but the sequence features that drive this association have not been fully explored. Here we train support vector machines (SVM) on sRNA sequencing data obtained from AGO-immunoprecipitation experiments to identify features that determine sRNA affinity to specific AGOs. Our SVM reveal that AGO affinity is strongly determined by complex k-mers in the 5’ and 3’ ends of sRNA, in addition to well-known features such as sRNA length and the base composition of the first nucleotide. Moreover, we find that these k-mers tend to overlap known transcription factor (TF) binding motifs, thus highlighting a close interplay between TF and sRNA-mediated transcriptional regulation. We embedded the learned SVM in a computational pipeline that can be used for de novo functional classification of sRNA sequences. This tool, called SAILS, is provided as a web portal accessible at http://sails.eu.nu.

## INTRODUCTION

The small RNA (sRNA) is a class of non-coding RNA with significant roles in developmental biology, physiology, pathogen interactions, and more recently in genome stability and transposable element control (1). Plants have two major classes of sRNA: the micro-RNA (miRNA), which is processed from imperfectly self-folded hairpin precursors derived from miRNA genes (2); and the small-interfering RNA (siRNA) that is produced from double-stranded RNA duplexes (3) (Fig. 1). siRNAs can be further divided into three major groups: secondary siRNA such as trans-acting (ta)-siRNAs, which are promoted by miRNA cleavage of messenger RNA; natural antisense transcript (nat)-siRNAs derived from the overlapping regions of antisense transcript pairs naturally present in the genome; and heterochromatin-associated (hc)-siRNAs, mostly generated from transposons, heterochromatic and repetitive genomic regions and involved in DNA methylation and heterochromatin formation.

Apart from their biogenesis, the mode of action of a given sRNA is tightly related with the Argonaute protein to which it can bind (4). Argonautes form the core of all sRNA-guided silencing complexes identified so far. Once loaded into Argonaute, sRNA guide the silencing machinery to targets through base pairing principles. Argonautes are highly conserved proteins with family members in most eukaryotes (4–6). Although there are two main subfamilies of Argonautes in eukaryotes: AGO and PIWI; only AGO proteins can be found in plants. Also in plants, AGOs can be grouped into three phylogenetic clades with a highly variable number of elements from species to species (5) (Fig. 2). Arabidopsis has ten members with specialized or redundant functions among them: AGO1, AGO5 and AGO10 in the first clade; AGO2, AGO3 and AGO7 form the second clade; and the third clade is composed by AGO4, AGO6, AGO8 and AGO9 (4). Members of the first and second clade are involved in post-transcriptional silencing (PTS) by inhibiting translation or by promoting messenger RNA cleavage, and AGOs in the third clade are chromatin modifiers that induce transcriptional silencing (TS) via mechanisms such as DNA methylation (7–9). Understanding the mechanisms that determine the loading of sRNA to specific AGOs is essential for predicting their biological function, and for identifying their putative silencing targets.

High throughput sequencing in combination with immunoprecipitation (IP) techniques have made possible to determine the sequences of sRNA that are bound to different AGO families. AGO-IP experiments have been performed for AGO1, AGO2, AGO4, AGO5, AGO6, AGO7, AGO9 and AGO10. The low expression level of AGOs 3 and 8 suggests that they may not be functionally relevant. Previous analyses have shown that AGO-sRNA associations are partly determined by the 5’ terminus and the length of a sRNA sequence (10, 11). Sequences of approximately 21 nucleotides (nt) tend to be involved in PTS, while 24 nt sRNAs are characteristic of TS. Although sequence length has been widely used as a way to infer the silencing pathway a given sRNA is most likely implicated in, by itself it is an inaccurate predictor since many sequencing products can lack other structural features known to enable AGO loading (10–13). Contrary to animals and flies, in plants the 5’ nucleotide is also recognized as a strong indicator of AGO sorting. Enrichment for sequences starting with pyrimidines are frequent in AGOs from the first clade (AGO1/10: uridine and AGO5: cytosine), adenosine dominates the third clade (AGO4/6/9) as well as AGO2 from the second clade. Furthermore, direct experimental evidence show that mutating the 5’ nucleotide of a sRNA can redirect its AGO destination (14); however, other relevant features, such as sequence motifs encoded by the primary structure appear to play a role (7, 15–18).

While sRNA that participate in PTS have been intensively studied, leading to the discovery of many structural features that influence activation, much less is known about AGO-associated hc-sRNA in transcriptional silencing. Indeed, most studies that use hc-sRNAs give a strong emphasis to sequence length ignoring other important aspects of the sRNA sequence that promote an active role in genomic regulation. Studying AGO-bound sRNA is a starting point to fill this gap and to improve our understanding on the relationship between the structure and function of sRNA in plants. Here we train support vector machines (SVM) on sRNA sequencing data obtained from AGO-IP experiments to identify features that determine sRNA affinity to specific AGOs. Our SVM reveal that AGO affinity is strongly determined by complex k-mers in the 5’ and 3’ ends of sRNA, in addition to well-known features such as sRNA length and the base composition of the first nucleotide. Moreover, we find that these k-mers tend to overlap known transcription factor (TF) binding motifs, thus highlighting a close interplay between TF and sRNA-mediated transcriptional regulation. We incorporated the learned SVM in an online computational pipeline that can be used for *de novo* sRNA functional classification. The classification pipeline is suitable for individual sRNA but also for high-throughput sRNA-seq datasets.

## MATERIAL AND METHODS

### Data sets

Table 1 summarizes the *A. thaliana* deep-sequencing sRNA libraries used in this study. For SVM model training and testing we used one Col-0 wild-type sRNA library as well as eight AGO-IP datasets. SVM model validation was afterwards performed on additional AGO-IP data from different tissues and from pathogen infected plants, in addition to a set of putative ta-siRNAs from several plant species. All sRNA-seq datasets were pre-processed and mapped to the Col-0 *A. thaliana* reference genome. Reads with at least one perfect match were collapsed into unique sRNA sequences and single copy sRNAs were removed. sRNA sequences in the genome-wide library that were not present in any of the AGO-IP sets, were isolated as a new group named “noAGO”. The “noAGO” set was used in the SVM training in combination with AGO-IP data to learn discriminative rules to identify sequences with low potential to load to an AGO and therefore with small chance of becoming functional sRNAs.

### Learning procedure

We developed a supervised machine learning approach rooted in the Support Vector Machine (SVM) algorithm to learn classifiers capable of determining AGO-sRNA affinity from the sRNA sequences alone (Supplementary Notes). Briefly, the complete inference system comprises 3 layers (Figure 3A): layer 1 includes a binary SVM model that filters out sequences that do not show strong evidence for binding to any of the known plant AGOs, and that therefore are expected to be inactive; layer 2 is composed by an ensemble of binary one-vs-one classifiers, each trained to explore the dissimilarities in AGO-bound sRNA sequences in a pairwise fashion; finally, layer 3 consists of a voting system that assigns a single score to each AGO using the decision values produced in the previous layer. Layers 2 and 3 are interconnected, since the outputs of the classifiers from the 2^nd^ layer serve as inputs to the 3^rd^ layer, and the 3^rd^ layer combines them to provide a more informative result. Layer 1 is independent of all other classifiers and can therefore be decoupled if desired. All sRNA sequences used in training, testing or validation were transformed into sets of features comprising:

i. Position specific base composition (PSBC) One way to convert a string into a numerical representation is by using flags for the presence of a given nucleotide in a determined sequence position. This is equivalent to mapping each sequence position to a four-dimensional feature space that represents each of the four possible bases in the DNA alphabet as follows:

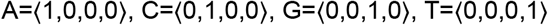 By using such a format, a sequence can be mapped to a feature space of dimensionality *F=A.L,* where *A* is the number of possibilities in the alphabet of nucleic acids and *L* expresses the dependence on the sequence length. Since the length of the sRNA sequences here analysed is not constant but can vary in an interval with an upper right limit here defined as being 27 bases, all sRNA must be projected into a space of size 27x4=108. In the case of sequences with a length shorter than 27, the extra positions in the feature vector can simply remain empty for all nucleotides. Although this approach is a reasonable solution to cope with the variation observed in length, it has the disadvantage of introducing noise in the representation that increases as the 3’ end of the model sequence is approached. This happens because the right most nucleotides in the real sRNA sequences are projected into more central positions of the model sequence, blurring 3’ side positional patterns when looking across instances with variable size. To compensate for that effect, the same kind of projection but starting at the 3’ position of the sRNA instead of the 5’ was additionally considered in a feature set here mentioned as PSBC2.
ii. k-mer composition Approaches based on k-mers map the presence or absence of subwords with a given length in the sRNA sequence into a feature space that represents all possible k-mers of that length. Taking as example k-mers of size 2, there are 4^2^=16 possibilities in the 4 letters universe of DNA. The 16 length 2-gram vector for the DNA ‘ACGT’ alphabet would then be:

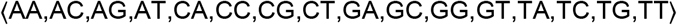 As an example, mapping the sequence “ATGCATG” onto this vector space, considering the presence or absence of each of the possible k-mers yields:

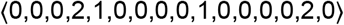 It is important to note that this method focuses on the frequency of patterns rather than their position in the sequence. Here k-mers of length 1 to 5 were explored.
iii. Shannon entropy scores Entropy as a measure of information content gives an indication about the degree of repetitiveness in a sequence. Among several flavours, Shannon entropy is one of the most popular and consists in a score given by:

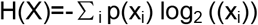

with *X* a sequence with length *l* and *p(x_i_)* the frequency of the character at position *i*.
iv. Sequence length This feature entails the number of nucleotides that compose each sRNA. Although simple, the enrichment for certain sizes in specific pathways is a consistent observation.

The learning methodology applied to layers 1 and 2 was similar. Prior to model training, highly correlated features (|Pearson score|>0.75) were removed keeping a single representative randomly selected from each correlated set. The remaining ones were normalized in the range between 0 and 1, to avoid dominance effects and numerical difficulties in the downstream calculations (19). SVM learning with recursive feature elimination (SVM-RFE) (20) was then applied to find models for layers 1 and 2 with a reduced and more informative feature set. To circumvent computational problems that typically arise when standard SVM algorithms are applied to large datasets, a linear kernel was employed with a specialized linear solver (21). A 5-fold cross-validation procedure was implemented to modulate data variation in each feature selection round and the mean ROC-AUC was calculated to assess the quality of the classifiers. Each round, 1/3 of the features with the lowest contribution for the discriminative model were eliminated, until 10 features were left. From here on, features were excluded one by one until no more features were available. The optimal feature subset for each classifier from layers 1 and 2 was determined with an elbow method applied to the curve formed by the mean cross-validation ROC-AUC values recorded during the feature selection process. The best features were subsequently used to train classifiers applying LIBSVM (22) with radial basis function (RBF) kernels to explore non-linear relationships in the data. A cascade scheme was implemented in this task to tackle computational problems that otherwise would not allow learning with such large datasets (Supplementary Notes, Figure 3B). To avoid biases in learning, training was performed with balanced datasets by under-sampling the largest class involved in each binary problem. Sequences with the highest read abundance were prioritized, and the remaining spots were occupied by instances randomly selected from the remaining pool(s).

In layer 3, a balanced dataset composed of sRNAs from all 8 AGO groups available was created. Decision values were computed for each of the sequences using the classifiers obtained for layer 2, and served as input for a 5-fold cross-validation procedure used to train and test the inference scheme applied to the 3^rd^ layer. Three strategies were explored to combine the outputs from layer 2: a voting system, where the winner is the AGO protein with the largest number of decisions in its favour; a weighted rule learned with a linear SVM algorithm; and a weighted rule learned with a non-linear SVM using a RBF kernel.

All SVM hyperparameters in layers 1, 2 and 3, were tuned by means of a grid search. A more detailed description of the SVM approach is provided in Supplementary Notes.

## RESULTS

### Detection of high confidence functional sRNAs

We explored a series of SVM classifiers to discriminate putatively functional from non-functional sRNA based on various sRNA sequence features. As outlined above (see section Material and Methods), SVM were trained on sequenced *A. thaliana* Columbia (Col-0) AGO-IP sRNA libraries in comparison with libraries of Col-0 total sRNA. Specific sRNA sequences contained in the AGO-IP sRNA libraries were labelled “AGO”, whereas sequences contained in the total library (but not in the AGO-sRNA libraries) were labelled “noAGO”. To minimize sequencing noise, single copy sRNA were removed.

### 5’ and 3’ k-mers are important in distinguishing functional from non-functional sRNA

We first trained separate SVM using either only the Position Specific Base Composition (PSBC or PSBC2) of the sRNA sequences, sRNA entropy, sRNA length, or all possible k-mers of size 1 to 5 nucleotides (see section Learning procedure). SVM trained on the PSBC or the k-mer features achieved considerable classification accuracy (~65%), while SVM trained using sRNA length and entropy performed less well (accuracy: ~50-60%, Figure 4). We then joint all the features together into a single classifier. Starting with 1582 features in total, our selection procedure yielded a final classifier containing only 138 features (Figure 4), and achieved a classification accuracy of nearly 80%. Of the 138 features, 127 are k-mers and 10 correspond to position specific nucleotides (Figure S2).

The k-mers of the final classifier were examined in more detail. Since the k-mers had no positional requirements for inclusion in the model we asked whether the k-mers retained in the final classifier showed any positional bias in the actual sRNA sequences. To do this, each of these k-mers was mapped back to the sRNA sequences contained in the “AGO” and the “noAGO” libraries. We found that the location of the k-mers was strongly biased toward the 5’ end of sRNA and to some extent also toward the 3’ end (Figure 5), with a visible depletion toward the centre of the sRNA sequence.

When looking to the 5’ end of sequences with meaningful length, we observed that 24 nt sRNA from the “AGO” set is dominated by k-mers starting with an adenine (A), and 21 nt sRNA have a mixture of adenine, uracil (U) and cytosine (C); plus in both cases there is absence of 5’ guanine (G), see Figure S4. This finding is consistent with previous reports showing that the base composition of the first nucleotide of a sRNA is important for AGO affinity (1, 4, 14). However, the vast majority (125 out of 138) of the features consisted of k-mers larger than one nucleotide in length, indicating that more complex sequence features guide AGO-sRNA associations.

### 5’ and 3’ k-mers are enriched for transcription factor binding motifs

We further assessed whether the k-mers correspond to known sequence motifs. To that end, we performed a motif analysis using the “AGO” and the “noAGO” sets with MEME (23), a computational framework for *de novo* and known motif identification. Then, we focused on k-mers with a size of 5 nt and mapped them to the motifs retrieved in the previous step. We found that the majority of k-mers (38 of 46) corresponded to segments of known or predicted motifs (Table S6), and noted a significant enrichment for k-mer matches in “AGO” when comparing with “noAGO” (100 and 32, respectively). Interestingly, the known motifs mapping k-mers were consistently related to stress response and development, which are processes known to be highly regulated by sRNAs (Table S6).

The TF enrichment was not surprising for 21nt sRNA which are known to act on genic sequences that frequently contain TF binding motifs. However, similar TF patterns were also found for 24nt sRNA, suggesting a role in gene regulation beyond the well-documented function of 24nt in heterochromatin silencing. We identified the 5-mer “AGAAG” as the k-mer that showed stronger enrichment in 24nt sequences compared with 21 nt sRNA from the “AGO” set and noted that this subsequence was associated with 3 motifs: At1g68670, a G2-like protein involved in phosphate homeostasis; SVP, a MADS protein that acts as a floral repressor and functions within the thermosensory pathway; and MYB77, which is expressed in response to potassium deprivation and auxin. All these motifs have been shown to have higher DNA-binding capacity when the targets are unmethylated (24), revealing a possible bridge between 24 nt heterochromatic sRNA, DNA methylation and TF occupancy.

### Classification and validation of specific AGO-sRNA associations

In the previous section our goal was to build a classifier that can distinguish functional from nonfunctional sRNA. We achieved this by training on “AGO” versus “noAGO” sRNA. A related problem is to find properties of sRNA that allow to infer their differential loading to specific AGO proteins, as this will determine their particular mode of action. To achieve this we trained binary one-vs-one classifiers to find sequence features that discriminate between the eight different Col-0 AGO-IP libraries (layer 2, Figure 3). The ensemble of one-vs-one classifiers was then subjected to a voting system that assigns a single score to each AGO (layer 3, Figure 3).

We used a 5-fold cross-validation scheme to determine the accuracy of the inference system at three levels: 1. AGO: fraction of sequences for which the AGO-bound protein was correctly predicted; 2. Clade: fraction of sequences for which the AGO prediction falls within the correct clade; and 3. Function: fraction of sequences for which the functional group can be correctly assigned based on the AGO prediction, translating predictions for AGO4/6/9 into a potential for involvement in TS, and assignments to other AGOs as suggestion of PTS activity. In addition to the 5-fold cross-validation scheme we also validated the classification system using 37 additional *A. thaliana* AGO-IP datasets that were never seen during the training phase by the classifier. These validation datasets were collected from different tissues and experimental conditions, which allowed us to evaluate the robustness of the classifier.

### Classification accuracy of sRNA at AGO, clade and function level

Results from our 5-fold cross-validation analysis showed that our classifiers achieved very high accuracy (Figure 6), indicating that differences between sRNA bound to specific AGOs can indeed be detected. Classification accuracy at the level of specific-AGO proteins was around 60% on average, ranging from as low as 40% (AGO7) to as high as 85% (AGO5), see Figure 6B. Since AGO proteins within a clade are highly homologous and similar in function, it is likely that sRNA-AGO binding is promiscuous in nature, which would render the search for discriminative features more challenging. Indeed, classification accuracy at the level of the clade and function were considerably higher than at the level of specific AGOs (clade accuracy: 85%, function accuracy 93%, see Figure 6A). Moreover, the final intra-clade classifiers were generally more complex than inter-clade classifiers, containing on average 122 features compared with 75 features, respectively (Figure 6C). Looking to the features retained in the final classifiers, intra-clade classifiers contained proportionally more k-mers of length larger than 1 nt compared with inter-clade classifiers (86% and 79% of the features in intra-clade and inter-clade models, respectively). Hence, factors that govern AGO-specific affinities within a clade appear to involve more complex sequence determinants. Similar to the AGO versus noAGO analysis presented above (section Detection of high confidence functional sRNAs), we found that the k-mers in the final classifiers showed strong positional bias toward the 5’ and 3’ ends of sRNA sequences (Supplementary Figure S3). This finding indicates that the same sRNA regions that differentiate functional from non-functional sequences also contribute to the binding affinity of sRNA to specific AGO proteins.

To study the nature of the information contained in the k-mers selected by SVM-RFE in more detail, we compared them with a motif analysis performed for each AGO library using the MEME suite (23). For a matter of simplicity, we focused on 5-mers (Supplementary Notes). We found that nearly 40% of the 5-mers were identified as motifs or derived segments (Table 2), often related to known transcription factors with roles in development and response to stress. For instance, in models involving AGO2 (an AGO linked to ta-siRNAs) we identified 5-mers matching the motif ETT, an experimentally validated target of an evolutionarily conserved ta-siRNA denominated tasiR-ARF (25). Targeting of ETT by ta-siRNA has been extensively studied and is known to have pleiotropic effects on Arabidopsis flower development by interference with the auxin pathway (26). These results thus support the conclusion that SVM learning captured sequence information with relevant biological meaning. A full overview of k-mers overlapping known or predicted motifs is provided in Table 2.

### Validation of the classification system

The AGO-sRNA classification framework was validated by testing on 35 AGO-IP libraries never seen during the training phase, including material acquired from different tissues and from plants under different treatment conditions (Table 1). These diverse datasets allowed us to evaluate the robustness of our classification framework. AGO, clade and function-based sensitivity were determined for each AGO library, in a procedure similar to the one applied to layer 3. Additionally, validation was also performed using two datasets of experimentally confirmed ta-siRNAs. One of these two datasets contained ta-siRNA only from *A. thaliana* while the other datasets comprised ta-siRNA from a large collection of different plant species. Since the ta-siRNA databases did not contain a record for the specific AGO to which the sequences load, function-based inference was the only adequate assessment in this case. Because no dataset of validated hc-siRNA is currently available, the quality of function inference for this kind of sRNA could not be measured.

Validation analysis revealed that our classifiers are relatively sensitive, even across biologically very heterogeneous datasets (Figure 7). Sensitivity was particularly high at the level of clade and function (clade: 70% and function: 84.3%), and moderate at the level of specific AGOs (AGO: 42%). Interestingly, when looking to the performance obtained for the ta-siRNA dataset, we observed high sensitivity values both for the set isolated for Arabidopsis (~80%) and also for the complete plant set (~95%). The sensitivity is considerably higher for the second set compared with the first which supports the idea that the inference system can identify PTS sRNAs not only from the species used in the learning phase but is also extensible to other plant species.

In conclusion, the inference system demonstrates robustness for tissue and treatment variation and can recognize sRNA membership independently of the cellular origin showing good generalization as desired for the discovery of new functional units.

## DISCUSSION

To our knowledge, this is the first time that classifiers were built to infer AGO-sRNA affinity from the sRNA sequence alone. Using adequate solvers and learning architectures, SVMs could be applied to large genomic datasets and discovered highly discriminative rules, both to distinguish sequences that bind to AGOs from other sequencing products, as well to infer AGO-sRNA kinship. In addition to the known 5’ nucleotide composition and the sequence length, feature selection revealed the contribution of other features, mostly complex k-mers, in defining sRNA preference for certain AGO proteins. The question how robust specific sRNA-AGO affinities are to changes in certain sequence properties is an interesting topic for future research and can shed light on evolutionary constraints on sRNA-mediated transcriptional and post-transcriptional silencing pathways. To answer such questions, additional experiments are necessary that can manipulate sRNA sequence in a precise and targeted fashion, for example by use of the CRISPR-Cas9 system. Although our inference method appear to be highly accurate in predicting the putative function of sRNA sequences, it is important to keep in mind that the actual biological activity of a given sRNA is dependent on other factors beyond the AGO loading step, such as the degree of complementarity between a sRNA and the target sequence, as well as the presence of specific chromatin states at the target locus.

AGO-sorting information has the potential to decrease the very high number of false positives reported by currently available PTS target prediction tools, but the true impact needs to be further evaluated. In any case, the computational framework here developed displays a discriminative power that makes it suitable for early screens in genome-wide sRNA libraries when looking for candidates with the highest chance to have certain functional roles, and when sorting sRNAs by TS and PTS classes is needed. The tool is ultimately a more affordable alternative to expensive and laborious AGO-IP experiments, since it can get AGO-sRNA profiles from a single genome-wide sequencing library. Another interesting application of the framework is to explore if specific sRNA from exogenous sources, such as artificially designed sequences or those derived from pathogens, correspond to functional plant sRNAs and which silencing pathways they are likely to target in a plant.

More sophisticated models can be developed using for example the expression patterns of the AGO proteins at the moment that the sRNA sequencing experiments take place. This extra information can eventually improve the capacity to correctly predict AGO affinity under particular situations, but on the other hand demands additional and more complex inputs, thus limiting the range of applications of the sRNA classifiers.

